# Fast myosin binding protein C knockout in skeletal muscle alters length-dependent activation and myofilament structure

**DOI:** 10.1101/2023.10.19.563160

**Authors:** Anthony L. Hessel, Michel Kuehn, Seong-Won Han, Weikang Ma, Thomas C. Irving, Brent A. Momb, Taejeong Song, Sakthivel Sadayappan, Wolfgang A. Linke, Bradley M. Palmer

## Abstract

In striated muscle, some sarcomere proteins regulate crossbridge cycling by varying the propensity of myosin heads to interact with actin. Myosin-binding protein C (MyBP-C) is bound to the myosin thick filament and is predicted to interact and stabilize myosin heads in a docked position against the thick filament and limit crossbridge formation, the so-called OFF state. Via an unknown mechanism, MyBP-C is thought to release heads into the so-called ON state, where they are more likely to form crossbridges. To study this proposed mechanism, we used the C2^-/-^ mouse line to knock down fast-isoform MyBP-C completely and total MyBP-C by ∼24%, and conducted mechanical functional studies in parallel with small-angle X-ray diffraction to evaluate the myofilament structure. We report that C2^−/−^ fibers presented deficits in force production and reduced calcium sensitivity. Structurally, passive C2^-/-^ fibers presented altered SL-independent and SL-dependent regulation of myosin head ON/OFF states, with a shift of myosin heads towards the ON state. Unexpectedly, at shorter sarcomere lengths, the thin filament was axially extended in C2^-/-^ vs. non-transgenic controls, which we postulate is due to increased low-level crossbridge formation arising from relatively more ON myosins in the passive muscle that elongates the thin filament. The downstream effect of increasing crossbridge formation in a passive muscle on contraction performance is not known. Such widespread structural changes to sarcomere proteins provide testable mechanisms to explain the etiology of debilitating MyBP-C-associated diseases.

## Main Text Introduction

The force-generating unit of striated muscle is the sarcomere and is predominately comprised of an interdigitating hexagonal array of thick and thin filaments (Fig.1A)^1,2^. Active force generation arises from the interaction of myosin heads projecting from the thick filament and actin in the thin filament, via so-called crossbridge cycling^3,4^. Separate from thin-filament-based calcium-dependent regulation of crossbridge formation by the troponin/tropomyosin^5^, the thick filament also plays an independent regulatory role in crossbridge formation^6–9^. In passive sarcomeres, each of the ∼300 myosin heads per thick filament exists in a conformational state on a spectrum between so-called “ON” and “OFF” states that affect crossbridge formation *during contraction*^10,11^. At one end is the OFF state, where the myosin head is docked against the helical tracks of the thick filament and has a reduced propensity to form a crossbridge upon activation. At the other end is the ON state, where the myosin head is positioned up and away from the thick filament, making it more likely to form a crossbridge upon activation^12^. Shifting a proportion of myosin heads toward the ON state increases the propensity to form crossbridges upon contraction^7,12–14^. Importantly, the sarcomere length (SL)-dependent transition of myosin heads toward the ON state with increasing SL is considered the mechanical underpinning of length-dependence of calcium sensitivity (length-dependent activation)^15,16^. However, myosin heads may also transition between ON and OFF states by SL-independent mechanisms, leading to a basal transition of myosin heads towards ON or OFF states while maintaining their SL-dependent property^8,10,17,18^.

**Figure 1.**
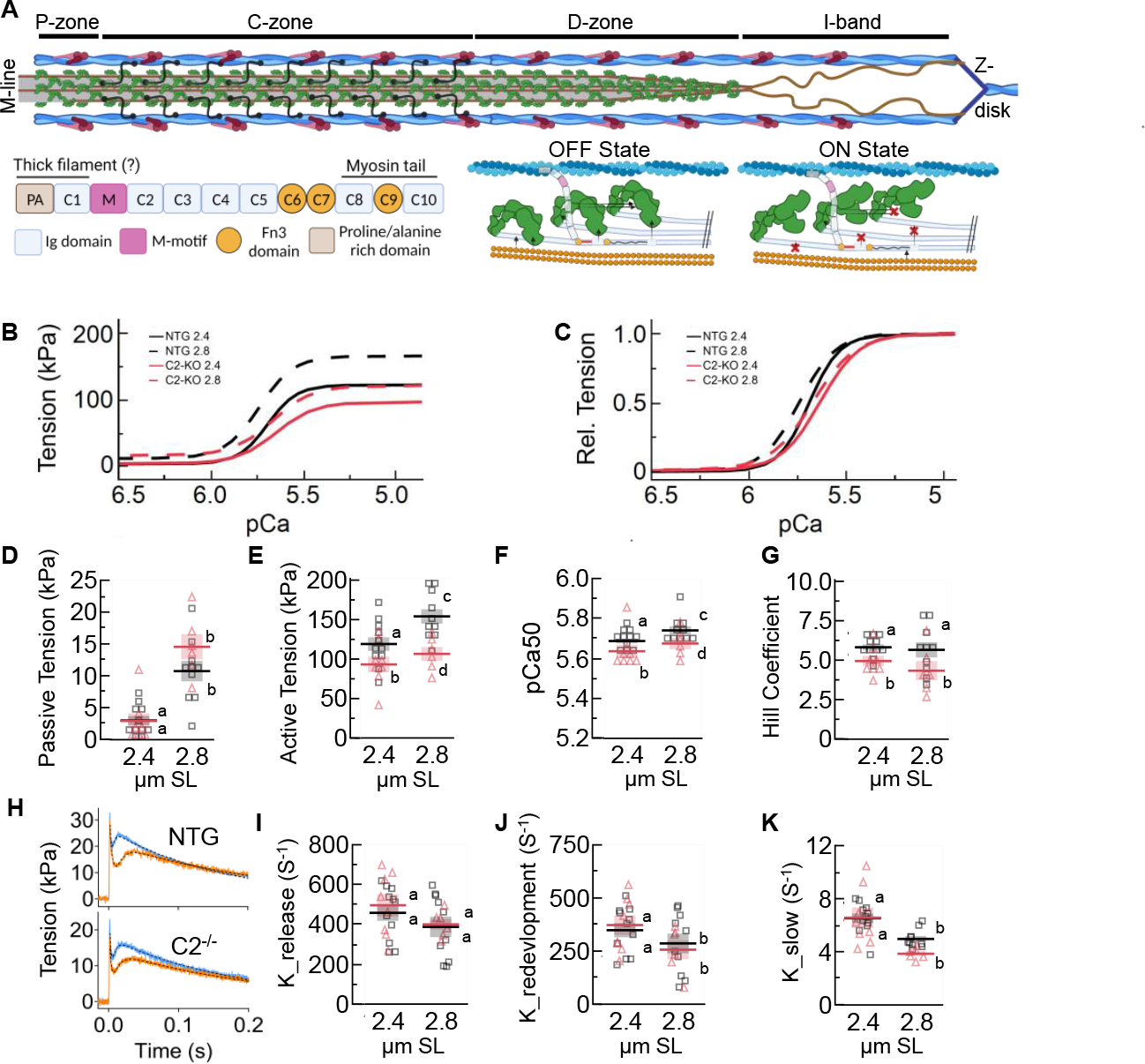
Mechanical assessment of permeabilized C2^-/-^ (black / squares) and NTG fibers (red / triangles) from EDL. (**A**) Cartoon representation of a half sarcomere. Thin filaments are comprised of actin filaments (blue), troponin-tropomyosin complexes (purple), and nebulin (not shown). Thick filament backbones (gray) are populated with myosin heads (green), titin filaments (brown), and MyBP-C (black). Thick filaments are demarcated into P-, C-, and D-zones, where MyBP-C is localized in the C-zone. In the I-band, titin extends from the Z-disk to the tops of the thick filament and produces titin-based force as an extensible spring. (**B**-**C**) Tension-pCa experiments for NTG (B) and C2^-/-^ (C) fiber bundles at 2.4 and 2.8 μm SL. (**D**) passive tension, (**E**) active tension, (**F**) pCa_50_, and (**G**) Hill coefficient were derived from tension-pCa experiments. Representative traces of quick-stretch – redevelopment experiments for NTG (top) and C2^-/-^ fibers (bottom) at 2.4 (orange and 2.8 (blue) μm SL. From these, the rate of force release (**I;** K_relase_) force redevelopment (**J;** K_redevelopment_) and slow phase (**K;** K_slow_) are calculated. Statistical results are presented as a connecting letters report, where different letters are statistically different (P < 0.05). Data reported as mean ± s.e.m. with full statistical details provided in Table 1.

Myosin-binding protein C (MyBP-C) is a proposed regulator of the myosin head ON/OFF state via stabilizing interactions between myosin heads in the OFF conformation^19–21^. MyBP-C arises at ∼43 nm intervals along the thick filament backbone, interactions with up to 108 myosin heads per thick filament^22,23^, with MyBP-C dysfunction associated with debilitating human myopathies^24–27^. Skeletal muscle MyBP-C is a chain of 10 domains (Fig. 1A) with the C’-terminus bound to the thick filament (C8-C10), and the other N’-terminal domains (C1-C7) pointed away from the thick filament, most likely interacting with myosin heads and the thin filament^22,23,28,29^. Skeletal muscles contain fast (fMyBP-C) and slow (sMpBP-C) isoforms that may function differently and are not necessarily fiber-type specific^19^. It was recently shown that rapid removal of the C1-C7 domains of fMyBP-C in the fast-twitch dominant psoas muscle led to an SL-independent movement of myosin heads towards the ON state, but the SL-dependent transition of myosin heads towards the ON state at longer vs. shorter SLs was largely intact^18^. These findings led to the hypothesis that the C1-C7 domains and C8-C10 domains regulate the SL-independent and SL-dependent components of myosin head ON/OFF regulation. If this is correct, then complete removal of MyBP-C could ablate, or at least reduce, both SL-independent and SL-dependent controls of myosin ON/OFF states. To explore the functional role of MyBP-C in skeletal muscle, we studied extensor digitorum longus (EDL) muscles from a fMyBP-C global knockout mouse (C2^−/−^) vs. age-matched non-transgenic (NTG) controls^20^. The NTG EDL is predominately a fast-twitch muscle with a ∼43% fMyBP-C expression^30^. C2^−/−^ express trace levels of fMyBP-C with a modest increase in sMyBP-C that leads to thick filaments with ∼24% fewer MyBP-C molecules^20^. We report that compared to NTG, C2^−/−^ fibers had reduced maximum tension and calcium sensitivity but retained length-dependence of calcium sensitivity. C2^−/−^ fibers presented an SL-independent shift toward the ON state, however, SL-length dependent structural changes were either altered or not detectable. Taken together, we provide evidence that MyBP-C plays a role in both the SL-dependent and SL-independent regulation of the myosin ON/OFF level in passive sarcomeres.

## Results and Discussion

We first evaluated the mechanical properties of permeabilized fiber bundles from C2^−/−^ and NTG EDL muscles. Tension-pCa measurements were made at SLs of 2.4 and 2.8 μm (Fig. 2B-C). In relaxed fibers (pCa 8), we observed the characteristic increase in passive tension from 2.4 to 2.8 μm SL but no detectable difference between genotypes (Fig. 1D; Table 1). During maximal contraction (pCa 4.5) both NTG and C2^−/−^ had increased tension at the longer SL, but C2^−/−^ produced less active tension across SLs (Fig. 1E; Table 1). NTG fibers increased pCa^50^ at the longer vs. shorter SL (Fig. 1F; Table 1), characteristic of a length-dependent activation^15,16^. While C2^−/−^ fibers also showed an increase in pCa^50^ at the longer SL, values were generally decreased across SLs (Fig. 1F; Table 1), as previously reported at 2.3 μm SL^20^, suggesting an SL-independent reduction in calcium sensitivity. No SL effect of the Hill coefficient was detected but there was a significant decrease for C2^−/−^ vs. NTG fibers (Fig. 1G; Table 1), which suggests a general SL-independent impairment to crossbridge recruitment in C2^−/−^ fibers. We next quantified force redevelopment after a small quick stretch that forcibly ruptured crossbridges for a measure of crossbridge kinetics (Fig. 1H). We found no genotype effects for the rate of force release (K_Release_), force redevelopment (K_Redevelopment_), or the slow phase (K_slow_) from the force redevelopment curve (see methods) were all decreased at the longer vs. shorter SL (Fig. 1I-K; Table 1), as expected^31^. Taken together, C2^−/−^ fibers present deficits in length-dependent enhancements normally observed with force production and calcium sensitivity, but length-dependence of crossbridge kinetics remains largely intact.

**Table 1.** Mechanical analysis from data in Figure 1. The ANOVA analysis F-stats and P-values are provided, as well as a connecting letter report from a Tukey HSD analysis. Data reported as mean ± s.e.m. *Significant (P < 0.05).

| Parameter | Genotype | SL ($\mu\text{m}$) | N | Mean | s.e.m. | Effect | F | P | Letters |
| --- | --- | --- | --- | --- | --- | --- | --- | --- | --- |
| K<sub>release</sub> (S<sup>-1</sup>) | WT | 2.4 | 11 | 458.09 | 40.12 | Genotype | 0.28 | 0.60 | a |
| K<sub>release</sub> (S<sup>-1</sup>) | WT | 2.8 | 10 | 387.68 | 51.53 | SL | 2.88 | 0.10 | a |
| K<sub>release</sub> (S<sup>-1</sup>) | C2<sup>-/-</sup> | 2.4 | 8 | 496.21 | 56.13 | Interaction | 0.06 | 0.81 | a |
| K<sub>release</sub> (S<sup>-1</sup>) | C2<sup>-/-</sup> | 2.8 | 6 | 400.06 | 26.91 |  |  |  | a |
| K<sub>red.</sub> (S<sup>-1</sup>) | WT | 2.4 | 11 | 348.46 | 33.33 | Genotype | 0.00 | 0.95 | a |
| K<sub>red.</sub> (S<sup>-1</sup>) | WT | 2.8 | 10 | 286.15 | 46.19 | SL | 4.17 | 0.04* | b |
| K<sub>red.</sub> (S<sup>-1</sup>) | C2<sup>-/-</sup> | 2.4 | 8 | 372.55 | 45.85 | Interaction | 0.37 | 0.55 | a |
| K<sub>red.</sub> (S<sup>-1</sup>) | C2<sup>-/-</sup> | 2.8 | 6 | 256.54 | 44.46 |  |  |  | b |
| K<sub>slow</sub> (S<sup>-1</sup>) | WT | 2.4 | 11 | 6.54 | 0.36 | Genotype | 6.23 | 0.02* | a |
| K<sub>slow</sub> (S<sup>-1</sup>) | WT | 2.8 | 10 | 4.98 | 0.22 | SL | 30.29 | <.0001* | b |
| K<sub>slow</sub> (S<sup>-1</sup>) | C2<sup>-/-</sup> | 2.4 | 8 | 6.58 | 0.78 | Interaction | 3.93 | 0.06 | a |
| K<sub>slow</sub> (S<sup>-1</sup>) | C2<sup>-/-</sup> | 2.8 | 6 | 3.86 | 0.18 |  |  |  | b |
| Min. Tension (mN) | WT | 2.4 | 11 | 2.93 | 0.72 | Genotype | 0.70 | 0.41 | a |
| Min. Tension (mN) | WT | 2.8 | 10 | 10.71 | 1.61 | SL | 48.56 | <.0001* | b |
| Min. Tension (mN) | C2<sup>-/-</sup> | 2.4 | 8 | 2.86 | 1.21 | Interaction | 1.16 | 0.29 | a |
| Min. Tension (mN) | C2<sup>-/-</sup> | 2.8 | 6 | 14.54 | 1.99 |  |  |  | b |
| Max. Tension (mN) | WT | 2.4 | 11 | 121.56 | 9.05 | Genotype | 11.28 | 0.002* | a |
| Max. Tension (mN) | WT | 2.8 | 10 | 164.21 | 9.76 | SL | 11.00 | 0.002* | c |
| Max. Tension (mN) | C2<sup>-/-</sup> | 2.4 | 8 | 95.80 | 10.61 | Interaction | 0.52 | 0.48 | b |
| Max. Tension (mN) | C2<sup>-/-</sup> | 2.8 | 6 | 120.75 | 9.66 |  |  |  | d |
| pCa<sub>50</sub> | WT | 2.4 | 11 | 5.69 | 0.01 | Genotype | 7.01 | 0.01* | a |
| pCa<sub>50</sub> | WT | 2.8 | 10 | 5.74 | 0.02 | SL | 4.28 | 0.04* | c |
| pCa<sub>50</sub> | C2<sup>-/-</sup> | 2.4 | 8 | 5.64 | 0.03 | Interaction | 0.07 | 0.79 | b |
| pCa<sub>50</sub> | C2<sup>-/-</sup> | 2.8 | 6 | 5.68 | 0.03 |  |  |  | d |
| Hill Coeff. | WT | 2.4 | 11 | 5.84 | 0.19 | Genotype | 6.91 | 0.01* | a |
| Hill Coeff. | WT | 2.8 | 10 | 5.68 | 0.48 | SL | 0.58 | 0.45 | a |
| Hill Coeff. | C2<sup>-/-</sup> | 2.4 | 8 | 4.97 | 0.33 | Interaction | 0.28 | 0.60 | b |
| Hill Coeff. | C2<sup>-/-</sup> | 2.8 | 6 | 4.36 | 0.61 |  |  |  | b |

**Figure 2.**
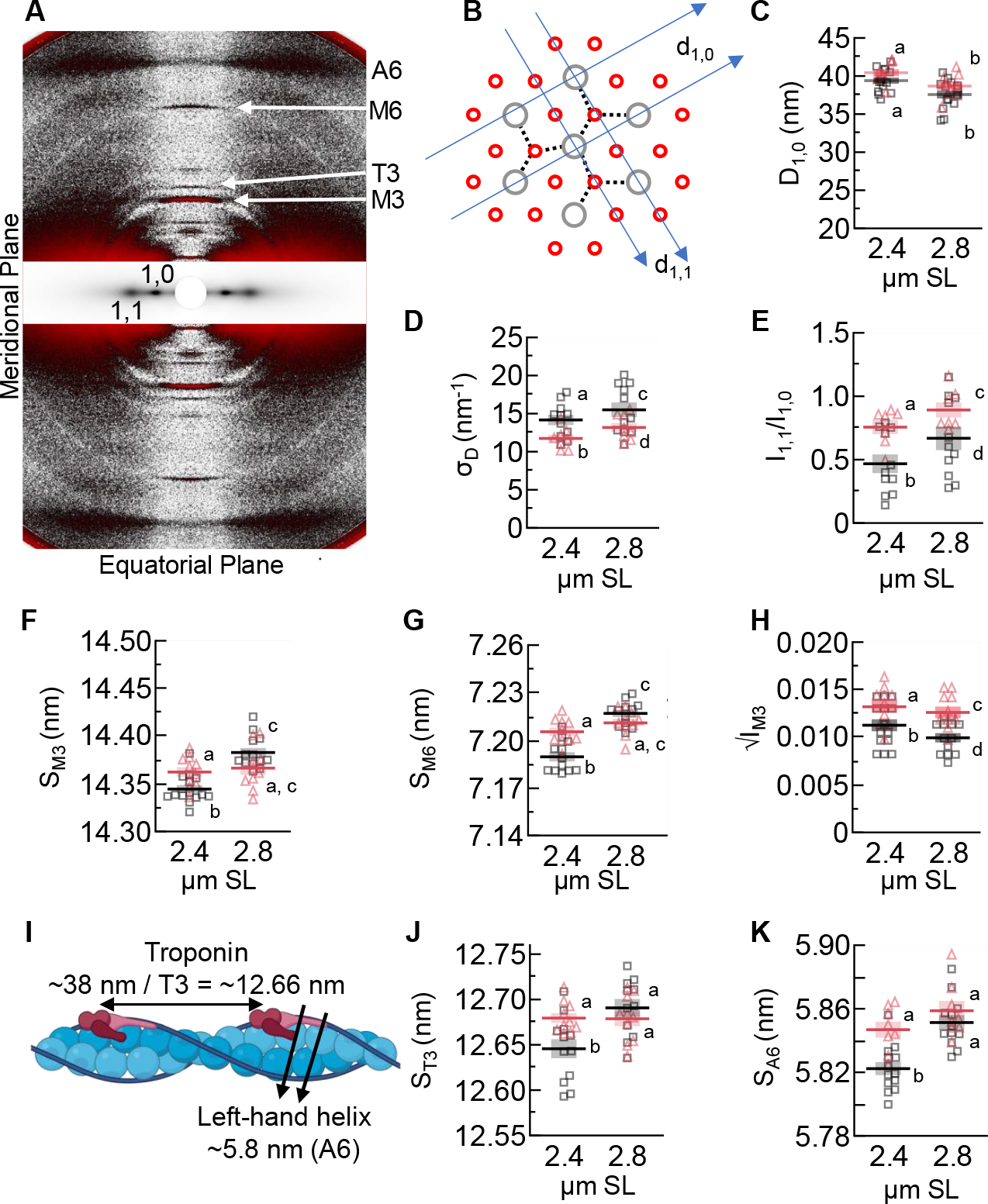
Sarcomere structures of C2^-/-^ (black / squares) and NTG fibers (red / triangles). (**A**) A representative image of an X-ray diffraction pattern, with reflections of interest labeled. The area around the equatorial axis was scaled differently to make reflections easier to view. (**B**) A cross-section of a myofibril in the thick (gray) and thin (red) filament overlap zone. Example myosin thick-thin filament crossbridges drawn (dotted lines). Overlayed are the geometric lattice planes d_1,0_ and d_1,1_, which lead to the 1,0 and 1,0 equatorial intensities, respectively. (**C**) d_10_ spacing quantifies lattice spacing. (**D**) σ_D_ quantifies lattice spacing heterogeneity. (**E**) I_1,1_ /I_1,0_ is a measure of mass distribution (i.e., myosin heads) between thick and thin filaments. (**F**) S_M3_ is the periodicity between myosin heads along the thick filament and indicates myosin head orientation. (**G**) S_M6_ is from a periodicity along the thick filament and quantifies the average thick filament length. (**H**) √IM3 is proportional to the electron density creating the reflection and can be interpreted as quantifying the orderness of myosin heads along the thick filament. (**I**) A cartoon representation of the thin filament, with periodicities of interest labeled. (**J**) S_T3_ is the (third-order) axial periodicity of troponin. (**K**) S_A6_ is the axial periodicity of the left-handed helix of actin and indicates thin filament twisting and elongation. Statistical results are presented as a connecting letters report, where different letters are statistically different (P < 0.05). Data reported as mean ± s.e.m. with full statistical details provided in Table 2.

We next evaluated myofilament structures using small-angle X-ray diffraction^12^. We collected X-ray diffraction patterns from relaxed (pCa 9) NTG and C2^-/-^ EDL fiber bundles at 2.4 and 2.8 μm SL. The myofilament lattice spacing was quantified via the 1,0 reflection (Fig. 2A) which represents the spacing of the d_1,0_ lattice plane within the filament overlap region (Fig. 2B). d_1,0_ decreased with increasing SL, as expected, with no genotype effect (Fig. 2C; Table 2). Lattice spacing heterogeneity (σ^D^)^32^ increased with increasing sarcomere length, as expected^33^, but was lower in C2^b^vs. NTG fibers across SLs (Fig. 2D; Table 2), suggesting that fMyBP-C impacts lattice order across SLs potentially via thin filament interactions^21,28^. sMyBP-C and fMyBP-C have well-characterized structural differences in their N-termini, yet differences in functionality are still not fully described^19,30^. With only sMyBP-C in the C2^-/-^ fibers and a mix of MyBP-C isoforms in the NTG, our results for d_1,0_ and σ_D_ suggest that sMyBP-C supports or allows a greater radial distance between myofilaments compared to fMyBP-C.

**Table 2.** Small angle X-ray diffraction analysis from data in Figure 2. The ANOVA analysis F-stats and P-values are provided, as well as a connecting letter report from a Tukey HSD analysis. Data reported as mean ± s.e.m. *Significant (P < 0.05).

| Parameter | Genotype | SL ($\mu\text{m}$) | N | Mean | s.e.m. | Effect | F | P | Letters |
| --- | --- | --- | --- | --- | --- | --- | --- | --- | --- |
| D<sub>1,0</sub> (nm) | WT | 2.4 | 14 | 39.39 | 0.41 | Genotype | 8.28 | 0.01* | a |
| D<sub>1,0</sub> (nm) | WT | 2.8 | 14 | 37.57 | 0.51 | SL | 97.33 | <.0001* | b |
| D<sub>1,0</sub> (nm) | C2<sup>-/-</sup> | 2.4 | 9 | 40.44 | 0.46 | Interaction | 11.57 | 0.0001* | a |
| D<sub>1,0</sub> (nm) | C2<sup>-/-</sup> | 2.8 | 9 | 38.68 | 0.54 |  |  |  | b |
| $\sigma_D$ (nm<sup>-1</sup>) | WT | 2.4 | 11 | 14.17 | 0.6866 | Genotype | 5.87 | 0.03* | a |
| $\sigma_D$ (nm<sup>-1</sup>) | WT | 2.8 | 11 | 15.50 | 0.9687 | SL | 22.27 | <.0001* | c |
| $\sigma_D$ (nm<sup>-1</sup>) | C2<sup>-/-</sup> | 2.4 | 8 | 11.74 | 0.4896 | Interaction | 0.02 | 0.98 | b |
| $\sigma_D$ (nm<sup>-1</sup>) | C2<sup>-/-</sup> | 2.8 | 8 | 13.18 | 0.5929 | | | | d |
| I<sub>1,1</sub>/I<sub>1,0</sub> | WT | 2.4 | 11 | 0.47 | 0.0735 | Genotype | 11.79 | 0.003* | a |
| I<sub>1,1</sub>/I<sub>1,0</sub> | WT | 2.8 | 11 | 0.67 | 0.0931 | SL | 3.54 | 0.04* | c |
| I<sub>1,1</sub>/I<sub>1,0</sub> | C2<sup>-/-</sup> | 2.4 | 7 | 0.76 | 0.0554 | Interaction | 1.13 | 0.34 | b |
| I<sub>1,1</sub>/I<sub>1,0</sub> | C2<sup>-/-</sup> | 2.8 | 7 | 0.89 | 0.0601 |  |  |  | d |
| S<sub>M3</sub> (nm) | WT | 2.4 | 14 | 14.344 | 0.0041 | Genotype | 5.17 | 0.03* | a |
| S<sub>M3</sub> (nm) | WT | 2.8 | 13 | 14.383 | 0.0047 | SL | 22.22 | <.0001* | c |
| S<sub>M3</sub> (nm) | C2<sup>-/-</sup> | 2.4 | 10 | 14.362 | 0.0051 | Interaction | 22.52 | <.0001* | b |
| S<sub>M3</sub> (nm) | C2<sup>-/-</sup> | 2.8 | 10 | 14.366 | 0.0071 |  |  |  | a, c |
| S<sub>M6</sub> (nm) | WT | 2.4 | 12 | 7.190 | 0.003 | Genotype | 0.01 | 0.91 | a |
| S<sub>M6</sub> (nm) | WT | 2.8 | 12 | 7.217 | 0.002 | SL | 58.77 | <.0001* | c |
| S<sub>M6</sub> (nm) | C2<sup>-/-</sup> | 2.4 | 10 | 7.206 | 0.003 | Interaction | 21.29 | <.0001* | b |
| S<sub>M6</sub> (nm) | C2<sup>-/-</sup> | 2.8 | 10 | 7.211 | 0.003 |  |  |  | a, c |
| $\sqrt{I_{M3}}$ | WT | 2.4 | 14 | 0.0113 | 0.0006 | Genotype | 9.64 | 0.01* | a |
| $\sqrt{I_{M3}}$ | WT | 2.8 | 13 | 0.0099 | 0.0005 | SL | 5.79 | 0.03* | c |
| $\sqrt{I_{M3}}$ | C2<sup>-/-</sup> | 2.4 | 10 | 0.0132 | 0.0006 | Interaction | 0.86 | 0.36 | b |
| $\sqrt{I_{M3}}$ | C2<sup>-/-</sup> | 2.8 | 10 | 0.0126 | 0.0006 | | | | d |
| S<sub>T3</sub> (nm) | WT | 2.4 | 12 | 12.645 | 0.0105 | Genotype | 0.96 | 0.34 | a |
| S<sub>T3</sub> (nm) | WT | 2.8 | 11 | 12.690 | 0.0098 | SL | 5.79 | 0.01* | c |
| S<sub>T3</sub> (nm) | C2<sup>-/-</sup> | 2.4 | 10 | 12.679 | 0.0054 | Interaction | 3.91 | 0.03* | b |
| S<sub>T3</sub> (nm) | C2<sup>-/-</sup> | 2.8 | 10 | 12.678 | 0.0082 |  |  |  | d |
| S<sub>A6</sub> (nm) | WT | 2.4 | 13 | 5.822 | 0.0041 | Genotype | 7.48 | 0.01* | a |
| S<sub>A6</sub> (nm) | WT | 2.8 | 12 | 5.851 | 0.0047 | SL | 14.20 | <.0001* | a |
| S<sub>A6</sub> (nm) | C2<sup>-/-</sup> | 2.4 | 8 | 5.847 | 0.005 | Interaction | 2.65 | 0.05* | b |
| S<sub>A6</sub> (nm) | C2<sup>-/-</sup> | 2.8 | 8 | 5.859 | 0.0062 |  |  |  | a |

The distribution of myosin heads between the ON and OFF states is a critical determinant of muscle performance during contraction (Fig. 2E)^8,11,15^. In mammalian muscle, there is a well-known SL-dependent mechanism by which myosin heads become ON: sarcomere stretch extends I-band titin, which increases the titin-based force that elongates the thick filament axially and ultimately leads to some myosin heads shifting from an OFF towards an ON state^12,33,34^. Elongating the thick filament most likely dissociates the stabilizing interactions between OFF myosin heads and the thick filament backbone (e.g. titin and MyBP-C)^22,23^. Other mechanisms regulate myosin heads in an SL-independent fashion, such as phosphorylation^35^ or the external addition of pharmaceuticals that force myosin heads into OFF or ON states^10,17,33^. Importantly, without SL change, repositioning myosin heads into the ON state still elongates the thick filament^10,17,33^, most likely due to structural rearrangements within the thick filament backbone that are not yet understood^11^. We tracked three X-ray reflections to study this phenomenon. 1) The intensity ratio between the 1,1 and 1,0 reflections (I_10_ / I_11_), which tracks the radial movement of mass in the form of myosin heads from thick toward neighboring thin filaments^36^. 2) The spacing and intensity of the M3 reflection (I_M3_, S_M3_). S_M3_ represents the average axial periodicity between myosin crowns along the thick filament. S_M3_ does not provide the radial position of the myosin heads off the thick filament backbone but does provide an orientation change that generally tracks the myosin ON/OFF state. Typically, increasing S_M3_ aligns with myosin head movement into an ON state, but is not strictly mandatory^13^. I_M3_ is an indicator of the helical ordering of the myosin heads and is a useful indicator of myosin structure change. Since the diffracted intensity is proportional to the square of the total electron density, the square root of I_M3_ (√I_M3_) is directly correlated to the number of diffracting myosin heads. All myosin heads can be in different orientations along the thick filament, so increasing or decreasing the homogeneity of myosin head positions will increase or decrease the I_M3_, respectively^37^. 3) The spacing of the M6 reflection (S_M6_) captures the coiled-coil periodicity of the myosin tail along the thick filament backbone and is used to measure thick filament elongation^38^.

For NTG fibers, we observed the typical SL-dependent increase of I_10_ / I_11_, S_M6_, S_M3,_ and √IM3 from the short to long SL (Fig. 2E-H; Table 2). Strikingly, C2^-/-^ fibers presented nearly constant values across SLs for S_M6_ and S_M3_. S_M6_ and S_M3_ values in C2^-/-^ fibers were generally elevated to the level of NTG fibers at the longer length, so that at the short SL, C2^-/-^ values were greater than NTG values (Fig. 2F-G; Table 2). These findings suggest two important conclusions. First, compared to NTG fibers, C2^-/-^ fibers have more myosin heads in the ON position across SLs. This can be caused by destabilization of the OFF state, which seems likely in the C-zone, as OFF-state myosin heads interact with MyBP-C domains C8-10^22,23^. Second, muscles with diseased, genetically modified, or partially cleaved MyBP-Cs all present evidence of destabilization of at least some C-zone myosin heads in the OFF state^18,20^. In theory, MyBP-C keeps a subpopulation of myosin heads in the C-zone in a different orientation than those in the P- or D-zones^37^. It seems reasonable that removing about 50% of the MyBP-C molecules from the thick filament in C2^-/-^ leads to those “freed” C-zone myosin heads assuming the orientation of their P-zone and D-zone counterparts, increasing overall myosin head order. Indeed, √I_M3_ was elevated in C2^-/-^ across SLs (Fig. 2H; Table 2), which indicates more myosin head order – something typically associated with myosins transitioning towards the OFF state, but in this case, where other markers indicate the opposite (see above), C-zone myosins being more ordered with D- and P-zone myosin heads seems more likely.

Surprisingly in C2^-/-^ fibers, SL-extension did not elongate the thick filament or reorient the myosin heads as is typical but did elevate the average position of myosin heads away from the backbone, as demonstrated by the increased I_10_ / I_11_, albeit with larger values across SL (Fig. 2E-H; Table 2). One possible explanation is that C2^-/-^ fibers were already transitioned to a higher ON state, even at short SL, and so the effect of stretch was reduced (and may have occurred below our detection limits). However, we could detect a radial head movement from short to long lengths in C2^-/-^ fibers, even though they were naturally transitioned to a more ON state across SLs. This may suggest that the radial movement of myosin heads toward the thin filament (I_11_/I_10_) is more sensitive to sarcomere length than orientation changes that alter myosin periodicity (S_M3_). It should be noted that the orientation of the blocked myosin head — the counterpart to the free head — also contributes to the M3 reflection and may also alter its typically OFF-state position in C2^-/-^ vs. NTG fibers. Taken together, the fMyBP-C KO in C2^-/-^ present altered SL-independent and SL-dependent regulation of myosin head ON/OFF states that is likely linked to an inability to stabilize the OFF state.

As a last assessment, we studied thin filament length. By a mysterious mechanism, thin filaments elongate with increasing SL in passive mammalian cardiac and skeletal muscle^18,33,39^ and we evaluated if this changes in fMyBP-C KO muscle. Evidence from direct visualization experiments^23,28,29^ shows so-called C-links, where the N’-terminal domains of MyBP-C interact with the thin filament, bridging the thick and thin filament. C-links, if present, would theoretically elongate the thin filaments during sarcomere stretch, but since C-links are short compared to an SL change (100’s of nm), the N’-terminal would drag along the surface of the thin filament. We quantified thin filament elongation by the spacing of the A6 reflection (S_A6_), which represents the periodicity of the left-handed helix of actin, and the T3 reflection (S_T3)_, which represents the 3^rd^-order axial spacing of troponin (Fig. H). In NTG fibers, we observed an increased S_A6_ and S_T3_ in the long vs. short SL (Fig. 2I-J; Table 2). In contrast, C2^-/-^ fibers presented no SL-dependence of S_A6_ or S_T3_ but had longer spacings at the short SL similar to those with longer SL (Fig. 2I-J; Table 2). While a loss of fMyBP-C and C-links could explain why there is little thin filament extension with increasing SL, it cannot explain why the thin filaments became longer at the short SL in C2^-/-^ vs. NTG fibers. In this study, C2^-/-^ sarcomeres have more ON myosin heads as well as longer thin filaments. In passive muscles, ON myosins produce a small number of crossbridges that generate a small amount of force on the thin filaments^40–43^. These bound crossbridges, estimated at ∼2% of myosin heads, increase with increasing proportion of ON myosin heads^44^. In C2^-/-^ fibers, more myosin heads are ON, and so we would predict more force-producing crossbridges as well, contributing to thin filament extension. These hypotheses could be tested by using mavacamten or dATP to force nearly all myosin heads into the OFF or ON state, respectively^10,45^, and tracking changes to thin filament length. The impact of this bridging and thin filament length on contraction performance is an unexplored area of muscle science.

## Materials and Methods

### Animal model and muscle preparation

Animal procedures were performed according to the *Guide for the Use and Care of Laboratory Animals* published by the National Institutes of Health and approved by the institutional animal care and use committee at the University of Vermont and the University of Cincinnati. Mice of either sex, 14-16 weeks old were deeply anesthetized with 2-4% isoflurane and killed by cervical dislocation. Skeletal muscles including extensor digitalis longus (EDL) were prepared as previously described^46^. Muscles were removed, and their tendons tied to wooden sticks to prevent contraction and placed in a relaxing solution composed of (mM): EGTA (5), MgCl_2_ (2.5), Na_2_H_2_ATP (2.5), imidazole (10), K-propionate (170), a protease inhibitor (1 minitab per 10 mL, Roche), pH = 7. Over the next 18 hours at 4°C, 50% of the relaxing solution was gradually replaced with glycerol. Samples were then stored at –20°C.

### Muscle Mechanics

Length-dependence of Ca^2+^sensitivity was assessed by performing force-pCa curves at sarcomere lengths of 2.4 (short) and 2.8 (long) μm using standard protocols^47^. Steady-state active force was assessed at pCa’s: 8.0, 6.33, 6.17, 6.0, 5.83, 5.67, 5.5, 5.0 (maximal [Ca^2+^]).

The solution contained (mM): EGTA (5), MgCl_2_ (1.12), BES (20), Na_2_H_2_ATP (5), Na-methyl sulfonate (67), PiOH (5), creatine phosphate (15), creatine kinase (300 U/mL), pH = 7. Force was normalized to the maximum force at pCa 5.0. A four-parameter Hill equation was fit to the normalized force-pCa data^48^ to calculate pCa_50_ (pCa at 50% maximum force; a measure of Ca^2+^ sensitivity) and the Hill coefficient. At pCa 5, a quick stretch of 0.25% muscle length was applied, and the force was fit to an equation of three exponentials: A exp (-k_release_ t) - B exp(-k_redevelopment_ t) + C exp(-k_slow_ t).

### Small angle X-ray diffraction and fiber mechanics apparatus

Samples were shipped to the BioCAT facility on ice for all experimental tests and stored at -20°C until used. On the day of experiments, EDL muscles were removed from the storage solution and vigorously washed in relaxing solution (composition (in mM): potassium propionate (45.3), N,N-Bis(2-hydroxyethyl)-2-aminoethanesulfonic acid BES (40); EGTA (10), MgCl_2_ (6.3), Na-ATP (6.1), DTT (10), protease inhibitors (complete), pH 7.0)). Whole fascicles were excised from EDL muscle and silk suture knots (sizing 6-0 or 4-0) were tied at the distal and proximal ends at the muscle-tendon junction as close to the fascicle as possible. Samples were then immediately transferred to the experimental chamber.

X-ray diffraction patterns were collected using the small-angle instrument on the BioCAT beamline 18ID at the Advanced Photon Source, Argonne National Laboratory^49^. The X-ray beam (0.103 nm wavelength) was focused to ∼0.06 x 0.15 mm at the detector plane. X-ray exposures were set at 1 s with an incident flux of ∼3x10^12^ photons per second. The sample-to-detector distance was set between 3.0 and 3.5 m, and the X-ray fiber diffraction patterns were collected with a CCD-based X-ray detector (Mar 165, Rayonix Inc, Evanston IL, USA). An inline camera built into the system allowed for initial alignment with the X-ray beam and continuous sample visualization during the experiment. Prepared fiber bundles were attached longitudinally to a force transducer (402A, Aurora Scientific, Aurora, Canada) and motor (322C, Aurora Scientific, Aurora, Canada), and placed into a bath of relaxing solution at 27°C. Force and length data were collected at 1000 Hz using a 600A: Real-Time Muscle Data Acquisition and Analysis System (Aurora Scientific, Aurora, Canada). Sarcomere length (SL) was measured via laser diffraction using a 4-mW Helium-Neon laser. The force baseline was set at slack length = 0 mN. After this initial setup, fiber length changes were accomplished through computer control of the motor, which we confirmed appropriate SL length change on a subset of samples. The mechanical rig was supported on a custom-designed motorized platform that allowed placement of muscle into the X-ray flight path and small movements to target X-ray exposure during experiments. Using the inline camera of the X-ray apparatus, the platform was moved to target the beam at different locations along the length of the sample. To limit X-ray exposure of any one part of the preparation, no part of the sample was exposed more than once.

### Experimental protocols and analysis

The experimental approach captured X-ray images in samples at two SLs across the *in vivo* physiological operating range^50^. Samples were stretched from 2.4 μm SL to 2.8 μm SL, at 0.1 μm SL s^-1^ with a 90 s hold phase to allow for stress relaxation. X-ray images were collected at the end of each hold phase.

### Analysis of X-ray diffraction patterns

X-ray images were analyzed using the MuscleX open-source data reduction package^51^. The “Quadrant Folding” routine was used to improve the signal-to-noise by adding together the four equatorial-meridional quadrants, which each provide the same information (Friedel’s Law). The “Equator” routine of MuscleX was used to calculate the I_1,1_ / I_1,0_ intensity ratio, d_1,0_ lattice spacing, and σ_D_. Meridional (M3, T3, M6) and off-meridional reflections (A6) were analyzed using the MuscleX “Projection Traces” routine. Spacing measurements of the meridional reflections were made in the reciprocal radial range ∼0 ≤ R ≤ 0.032 nm^-1^ for M3, M6, and T3 reflections, and ∼0.013 ≤ R ≤ 0.053 nm^-1^ for the A6 reflection, where R denotes the radial coordinate in reciprocal space^52^. Every image provides intensities of different quality, which leads to various levels of Gaussian fit errors for each intensity modeled, which increases the variation in spacings in the dataset. To limit these effects, fit errors > 10% were discarded. Positions of X-ray reflections on the diffraction patterns in pixels were converted to sample periodicities in nm using the 100-diffraction ring of silver behenate at d_001_ = 5.8380 nm. Intensity was normalized by the radially symmetric background measured by the “Quadrant Folding” routine.

### Statistics

Statistical analysis was conducted using JMP Pro (V16, SAS Institute, USA). The significance level was always set at α = 0.05. We used a repeated-measures analysis of variance (ANOVA) design with fixed effects SL, genotype, SL x genotype interaction term, and a nested random (repeated-measures) effect of the individual (when appropriate). Data was best Box-Cox transformed to meet assumptions of normality and homoscedasticity when necessary, which were assessed by residual analysis, Shapiro-Wilk’s test for normality, and Levene’s test for unequal variance. Significant main effects were subject to Tukey’s highly significant difference (HSD) multiple comparison procedures to assess differences between factor levels. This data is indicated in graphs via so-called connecting letters, where factor levels sharing a common letter are not significantly different from each other. All data presented as mean ± s.e.m.

## Data availability statement

Datasets used to generate the figures and tables are included in supplemental information. Additional data that support the findings of this study are available from the corresponding authors upon reasonable request.

## Acknowledgments

We thank the BioCAT beamline support staff at the APS for their steadfast commitment to our project, and Anna Good for critical text and artistic editing. Some of the figures here were designed using the program Biorender. This research used resources of the Advanced Photon Source, a U.S. Department of Energy (DOE) Office of Science User Facility operated for the DOE Office of Science by Argonne National Laboratory under Contract No. DE - AC02 - 06CH11357, and further NIH support. The content is solely the responsibility of the authors and does not necessarily reflect the official views of the National Institute of General Medical Sciences or the National Institutes of Health.

## Funding

German Research Foundation grant 454867250 (ALH)

German Research Foundation grant SFB1002,A08 (WAL)

IZKF Münster Li1/029/20 (WAL)

National Institutes of Health P41 GM103622(TI), P30 GM138395 (TI), R01 AR079435 (SS), R01 AR079477 (SS), R01 HL130356 (SS), R01 HL105826 (SS), R01 AR078001 (SS), and R01

HL143490 (SS)

The American Heart Association 19UFEL34380251 (SS), 19TPA34830084 (SS) and 945748 (SS)

The PLN Foundation (SS)

## Author Contributions

Conceptualization: BMP, ALH Methodology: ALH, BMP,

BAM, TS, SS Investigation: BMP, ALH, MK, SH,

WM Visualization: ALH, BMP, WM, WL, TCI

Funding acquisition: BMP, ALH, TCI, TS, SS, WL

Project administration: ALH, BMP

Supervision: ALH, BMP Writing –

original draft: ALH

Writing – review &

editing: All authors

## Materials and Correspondence

Materials and correspondence requests can be submitted to Dr. Anthony Hessel and Dr. Bradley Palmer.

